# A preferential role for ventromedial prefrontal cortex in assessing “the value of the whole” in multi-attribute object evaluation

**DOI:** 10.1101/2020.09.29.319293

**Authors:** Gabriel Pelletier, Nadav Aridan, Lesley K. Fellows, Tom Schonberg

## Abstract

Everyday decision-making commonly involves assigning values to complex objects with multiple value-relevant attributes. Drawing on object recognition theories, we hypothesized two routes to multi-attribute evaluation: assessing the value of the whole object based on holistic attribute configuration or summing individual attribute-values. In two samples of healthy human participants undergoing eye-tracking and fMRI while evaluating novel pseudo-objects, we found evidence for both forms of evaluation. Fixations to, and transitions between attributes differed systematically when value of pseudo-objects was associated with individual attributes or attribute configurations. Ventromedial prefrontal cortex (vmPFC) and perirhinal cortex were engaged when configural processing was required. These results converge with our recent findings that individuals with vmPFC lesions were impaired in decisions requiring configural evaluation, but not when evaluating “the sum of the parts”. This suggests that multi-attribute decision-making engages distinct evaluation mechanisms relying on partially dissociable neural substrates, depending on the relationship between attributes and value.

**SIGNIFICANCE STATEMENT:** Decision neuroscience has only recently begun to address how multiple choice-relevant attributes are brought together during evaluation and choice among complex options. Object recognition research makes a crucial distinction between individual attribute and holistic/configural object processing, but how the brain evaluates attributes and whole objects remains unclear. Using fMRI and eye-tracking, we found that the vmPFC and the perirhinal cortex contribute to value estimation specifically when it emerged from the whole objects i.e. predicted by the unique configuration of attributes, and not when value was predicted by the sum of individual attributes. This novel perspective on the interactions between subjective value and object processing mechanisms bridges an important gap between the fields of object recognition and reward-guided decision-making.

## INTRODUCTION

Choosing which snack to buy requires assessing the value of options based on multiple attributes (e.g. color, taste, healthiness). Value can be related to individual attributes: for example, if someone loves chocolate, all snacks containing this ingredient will be valued above those that do not. Value can also emerge from the combination of individual attributes, such as for chocolate-peanut snacks, where the combination of sweet and salty ingredients within the same snack might yield a value greater than the sum of the individual attributes.

The object processing literature has shown that there are distinct neural substrates representing the individual elements that make up complex objects and the holistic, configural combinations of those elements, hierarchically organized along the ventral visual stream (VVS) (Riesenhuber and Poggio, 1999; Bussey and Saksida, 2002). Lesions to the perirhinal cortex (PRC), a medial temporal lobe structure situated at the anterior end of the VVS, impair object discrimination based on attribute configuration but spare discrimination based on individual attributes (Bussey et al., 2005; Bartko et al., 2007; Murray et al., 2007). Neuroimaging studies have shown that BOLD fMRI and PET signals in human PRC is more sensitive to multi-attribute configuration than to the component attributes of objects, whereas the lateral occipital cortex demonstrates higher sensitivity to single attributes compared to anterior regions of the VVS (Devlin and Price, 2007; Erez et al., 2016). This suggests that configural object recognition is supported by the PRC, and that individual attribute representations at earlier stages of object processing are sufficient for object recognition or discrimination under certain conditions.

Leading neuroeconomic models suggest that the ventromedial prefrontal cortex (vmPFC) encodes subjective value across stimuli as a “common currency” to support flexible decision-making (Chib et al., 2009; Levy and Glimcher, 2012; Delgado et al., 2016). While many of these studies presented multi-attribute objects (e.g. foods, trinkets), they have only rarely considered how the values of multiple attributes are combined. A handful of fMRI studies examined the neural correlates of options explicitly composed of multiple attributes. These have found that signal within the vmPFC reflect the integrated value of the component attributes when each independently contributes to value, i.e. when value is associated with individual elements of the option (Basten et al., 2010; Philiastides et al., 2010; Kahnt et al., 2011; Park et al., 2011; Lim et al., 2013; Hunt et al., 2014; Suzuki et al., 2017; Kurtz-David et al., 2019). However, these studies did not address whether there are distinctions in the neural processes underlying value construction based on summing attributes versus value emerging from the holistic configuration of attributes.

Recent evidence argues that the distinction between configural and elemental processing is important in valuation, just as it is known to be important in complex object recognition. We recently found that lesions to the vmPFC in humans impaired decisions between objects when value was associated with the configural arrangement of attributes, but spared decisions when value was associated with individual attributes (Pelletier and Fellows, 2019). Here, we employ a triangulation approach (Munafò and Smith, 2018) to further test this hypothesis using fMRI and eye-tracking to examine the neural and behavioural correlates of multi-attribute valuation in healthy women and men.

We hypothesized that estimating the value of multi-attribute visual objects in a condition where value was predicted by attribute configuration would engage the vmPFC as well as regions involved in complex object recognition (i.e. PRC) to a greater extent than an elemental condition where individual attributes contributed independently to the overall object value. We further hypothesized that fixations to, and fixation transitions between value-predictive attributes would differ between configural and elemental value conditions. We report data from two independent samples of healthy participants: one behavioural and eye-tracking study, and another that also included fMRI. An additional pilot study was carried out to determine the fMRI study sample size. All hypotheses and analysis steps were preregistered (osf.io/4d2yr).

## METHODS

Data were collected from three independent samples using the same experimental paradigm. This paradigm involved first learning and then reporting the monetary value of novel, multi-attribute pseudo-objects under elemental or configural conditions. We collected an initial behavioural sample to characterize learning, decision-making and eye gaze patterns. We then undertook a pilot fMRI study to estimate the sample size needed to detect effects of interest. Informed by this pilot study, a third sample underwent fMRI and eye-tracking. Data from the behavioural sample informed the preregistration of eye-tracking hypotheses to be replicated in the fMRI sample.

### Participants

Participants were recruited from the Tel Aviv University community via online advertising and through the Strauss Imaging Center’s participant database. Participants were healthy volunteers, with normal or corrected-to-normal vision, without any history of psychiatric, neurological, or metabolic diagnoses, and not currently taking psychoactive medication. The study was approved by the ethics committee at Tel Aviv University and the institutional review board of the Sheba Tel-Hashomer Medical Center.

#### Behavioural study

Forty-two participants were recruited to take part in the behavioural experiment. Nine participants were excluded due to poor task performance according to the exclusion criteria detailed below. The final behavioural sample included 33 participants (15 women, mean age 22 y, range 18-32). Eye tracking data were not available for three participants due to poor calibration of the eye-tracker.

#### fMRI pilot study

Imaging data were collected in a pilot sample of 8 participants (four women, mean age 25 y, range 21-31) to calculate the sample size needed to detect a significantly stronger modulation of value in the configural compared to the elemental trials in the vmPFC at an alpha level of 0.05 with 95% power. Power calculations were carried out with the fmripower software (http://fmripower.org/)(Mumford and Nichols, 2008), averaging beta weights for the contrast of interest across all voxels of a pre-defined brain region. Based on these calculations, we preregistered that 42 participants would be required. This sample size was also sufficient to detect a significant effect for the parametric modulation of value in the Configural condition alone, in the vmPFC (38 participants needed for 95% power). The vmPFC region of interest and the model used to analyse the pilot data are described below. Imaging data used for power and sample-size calculations are available on OpenNeuro (https://openneuro.org/datasets/ds002079/versions/1.0.0), and the code used to create the power curves and the vmPFC ROI mask are available with the preregistration document (osf.io/4d2yr). Pilot participants were not included in the final sample.

#### fMRI study

Fifty-five participants were recruited to take part in the full fMRI experiment. Nine participants were excluded due to poor task performance in the scanner, according to the preregistered exclusion criteria. Three participants were excluded because of MR artefacts, and one participant was excluded due to excessive motion inside the scanner based on fMRIprep outputs (Esteban et al., 2019). The final fMRI sample thus included 42 participants (21 women, mean age 27 y, range 18-39). Eye-tracking data could not be collected in 9 participants due to reflections caused by MR-compatible vision correction glasses.

### Experimental paradigm

The experimental paradigm was adapted from a recently published study (Pelletier and Fellows, 2019). Participants learned the monetary values of novel multi-attribute pseudo-objects (fribbles) in two conditions (configural and elemental), after which they were scanned while bidding monetary amounts for the objects. Fribbles were developed to study object recognition, and are designed to mimic real-world objects (Williams, 1998). They are composed of a main body and four appendages which we refer to as attributes, each available in three variations. Two fribble sets were used, one for each condition (randomly assigned for each participant); each set had the same body but different appendages.

In the configural condition, value was associated with the unique configuration (conjunction) of two attributes. In the elemental condition, value was associated with each of two individual attributes, which then could be combined to obtain the value of the whole object. Four different object sets were used across participants; the object set-condition assignment was counterbalanced. Learning order was counterbalanced across participants (configural followed by elemental or vice versa) and the order of object presentation was randomized in all experiment phases. An example of the stimuli as well as the value associations are shown on **Figure 1**.

**Figure 1.**
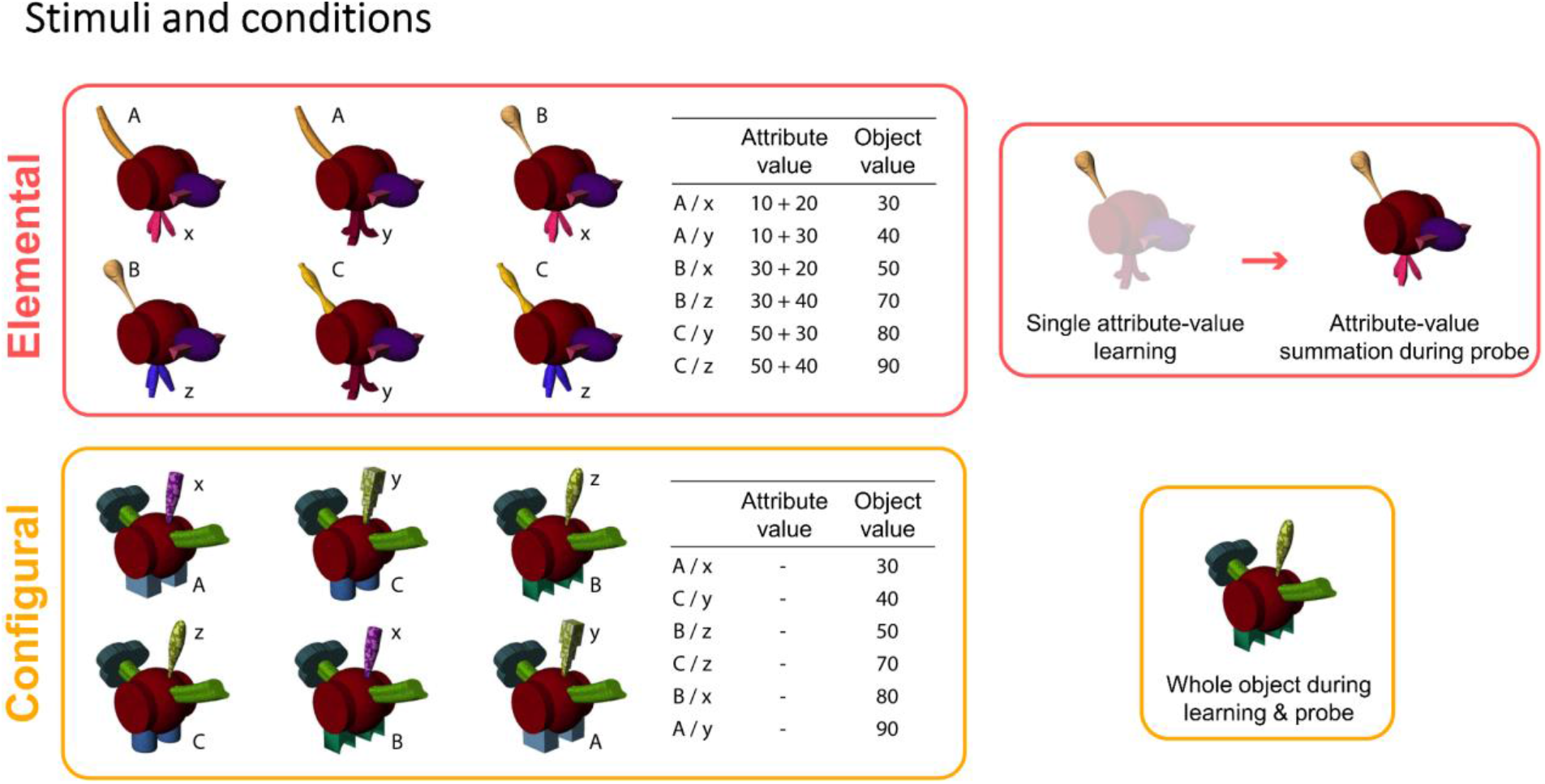
Stimuli and conditions. Stimuli and conditions. Example of fribble sets and object-average value associations. In the elemental condition, each fribble presented in the bidding phase had two individually value-predictive attributes which could be summed to estimate the value of the whole object. Objects were masked during the learning blocks so that value was clearly associated with a single salient attribute. In the learning probe and bidding phase, the unmasked objects were shown and object value was assessed by summing the individual attribute values (top right). In the configural condition, each fribble had two attributes which reliably predicted value only when appearing together, i.e. in configuration.

#### Learning phase

Participants were instructed before the experiment that they were acting as business owners, buying and selling novel objects. Before acquiring objects in their own inventory, they began by observing objects being sold at auction to learn their market price.

The learning phase included five learning blocks and one learning probe per condition. A block began with a study slide displaying all 6 objects to be learned in that condition, along with the average value of each object, giving the participant the opportunity to study the set for 60 s before the learning trials (**Fig. 2*A***). The learning trials began with the presentation of an object in the center of the screen above a rating scale, asking “How much is this item worth?”. Participants had 5 s to provide a value estimate for the object, using the ‘left’ and ‘right’ arrow keys to move a continuous slider and the ‘down’ arrow key to confirm their response. Feedback was then provided indicating the actual selling price of the object, with a bright yellow bar and the corresponding numerical value overlaid on the same rating scale. The object, rating slider and feedback were displayed for 2 s, followed by 2 s fixation cross. Each learning block presented all 6 objects 6 times each in random order for a total of 36 trials. After five learning blocks, learning was assessed with a probe consisting of 24 trials of the 6 learned objects presented four times each, in random order. The structure of probe trials was identical to the learning trials, but no feedback was given after the value rating.

**Figure 2.**
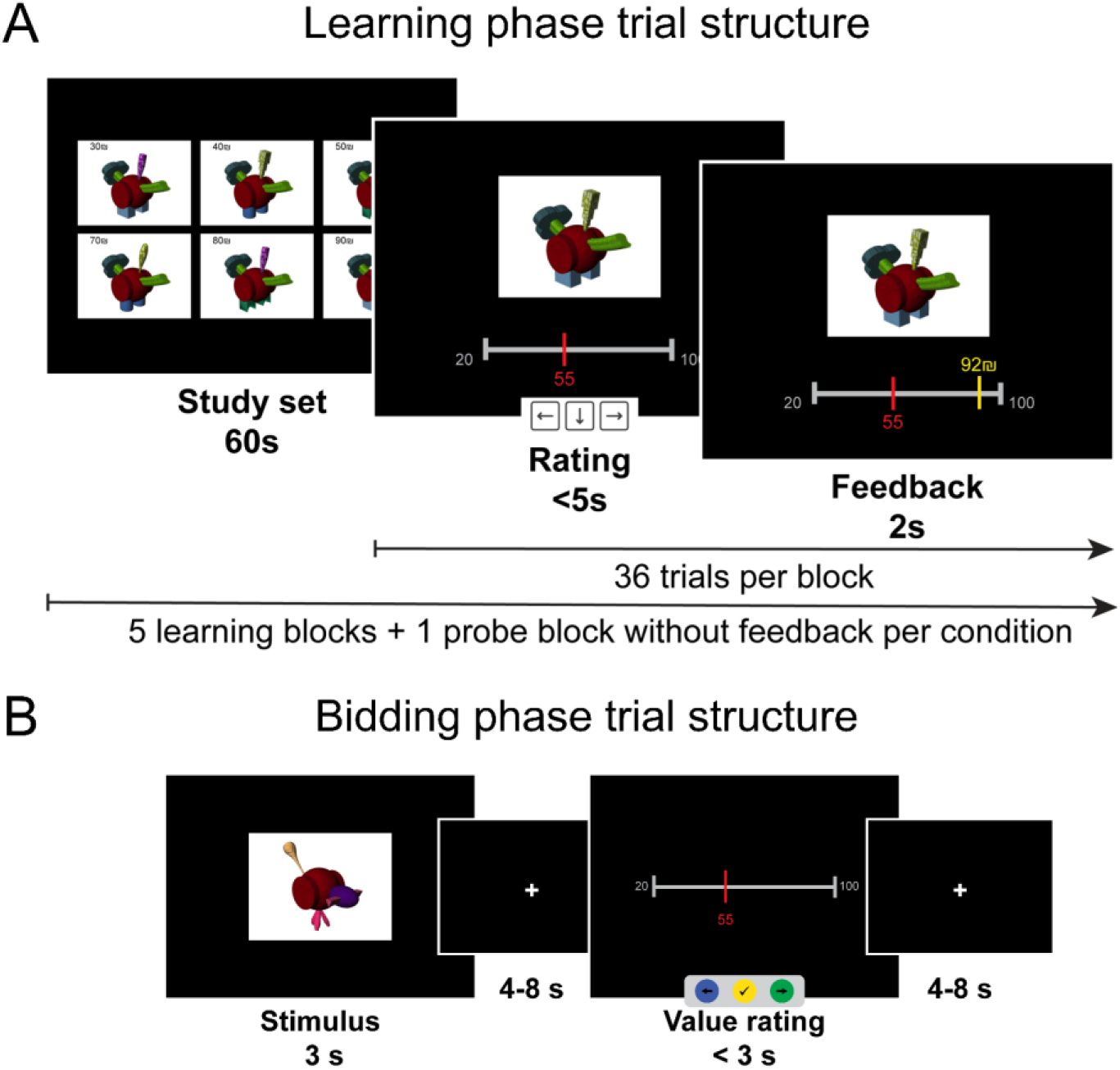
Experimental paradigm. **A)** Structure of a learning block. **B)** Trial structure of the bidding (fMRI) task.

In the elemental condition, values were associated with individual attributes. During the learning blocks, the object’s body and irrelevant attributes were occluded with a 50% transparent white mask, making the specific value-predictive attribute more salient (**Fig. 1**). Participants were told that value was associated only with the unmasked attribute. During the learning probe, objects were presented without masks, so all attributes were equally salient, and participants were instructed to sum the values of the two attributes they had learned.

In the configural condition, objects were displayed without masks during the entire learning phase, and the value of the object was associated with the unique configuration of two attributes. In this condition, participants could not learn object-values by associating value with any single attribute, because each attribute was included in both a relatively high-value and a relatively low-value object, as depicted in the object-value table (**Fig. 1**).

After learning, each of the 6 objects of the elemental condition had the same overall-value (sum of the two attribute-values) as one of the 6 configural objects. The object set in each condition contained 6 value-relevant attributes, each of which was part of two different objects in each set.

#### Bidding task

After learning, participants placed monetary bids on the learned objects to acquire them for their inventory while eye movements were tracked and, in the fMRI studies, fMRI was acquired. The task comprised four runs (scans) each containing the 12 objects (6 per condition) repeated twice in random order for a total of 24 trials. The structure of a bidding trial is depicted in **Fig. 2*B***. Before the bidding task, participants performed one practice run to familiarize themselves with task timings.

To make the task incentive-compatible, participants were instructed beforehand that all auctions would be resolved at the end of the session. If they bid sufficiently close to, or higher, than the true (instructed) object’s value, this object would be acquired and placed in their inventory. After the task, we would buy all the items in their inventory with a profit margin for the participant (similar to the situation where stores sell their products for a higher price than they paid from the manufacturer). The additional bonus compensation was calculated by summing the total amount paid by the experimenter to buy the participant’s inventory, minus the total of the bids placed by the participant to acquire these items. The margins were set so that the maximum bonus could not exceed 10 ILS (~$3 USD equivalent). Participants were told that they could not lose money in the experiment; if the total of their bids was substantially higher than the total retail value of their inventory, the bonus compensation was 0.

#### Anatomical scans and functional localizer task

After the bidding task, FLAIR and T1 anatomical scans and B0 field maps were acquired for the fMRI samples, with the parameters detailed below. Following structural scan acquisition, participants performed a functional localizer task adapted from (Watson et al., 2012) to define participant-specific visual regions of interest for analysis of the bidding task. Images from four categories (faces, scenes, objects and scrambled objects) were presented in blocks of 15 s, each containing 20 images displayed for 300 ms with a 450 ms inter-stimulus interval. Participants were instructed to press a button using the index finger of the right hand when an image was repeated twice in a row (1-back). The task was comprised of 4 runs of 12 blocks each. A 15 s fixation block ended each run. One run contained three blocks of each image category in a counterbalanced order.

### Data acquisition

#### Behavioural data

All phases of the experiment were programmed in Matlab (R2017b, The Mathworks, Inc.), using the Psychtoolbox extension (PTB-3) (Brainard, 1997). During the learning phase, and during the bidding task for the behavioural sample, stimuli were displayed on a 21.5-inch monitor and responses were made using a standard keyboard. We recorded value rating and reaction time for each learning trial.

During the bidding task in the fMRI, stimuli were presented on a NordicNeuroLab 32” LCD display (1,920 × 1,080 pixels resolution, 120 Hz image refresh rate) that participants viewed through a mirror placed on the head coil. Participants responded using an MR-compatible response box. Value rating, reaction time, and the entire path of the rating slider were recorded for each trial.

#### Eye tracking data

We recorded eye gaze data during the bidding task using the Eyelink 1000 Plus (SR research Ltd., Kanata, Ontario, Canada), sampled at 500 Hz. Nine-point calibration and validation were carried out before each run of the task.

#### fMRI data

Imaging data were acquired using a 3T Siemens Prisma MRI scanner and a 64-channel head coil. High-resolution T1-weighted structural images were acquired for anatomical localization using a magnetization prepared rapid gradient echo (MPRAGE) pulse sequence (Repetition time (TR) = 2.53 s, echo time (TE) = 2.99 ms, flip angle (FA) = 7°, field of view (FOV) = 224 × 224 × 176 mm, resolution = 1 × 1 × 1 mm).

Functional imaging data were acquired with a T2* weighted multiband echo planar imaging protocol (TR = 1,200 ms, TE = 30 ms, FA = 70 degrees, multiband acceleration factor of 4 and parallel imaging factor iPAT of 2, scanned in an interleaved fashion). Image resolution was 2 × 2 × 2 mm voxels (no gap between axial slices), FOV = 97 × 115 × 78 mm (112 × 112 × 76 acquisition matrix). All images were acquired at a 30° angle off the anterior–posterior commissures (AC–PC) line, to reduce signal dropout in the ventral frontal cortex (Deichmann et al., 2003). We also calculated field maps (b0) using the phase encoding polarity (PEPOLAR) technique, acquiring three images in two opposite phase encoding directions (anterior-posterior and posterior-anterior), to correct for susceptibility induced distortions.

### Data and code accessibility

Unthresholded whole-brain statistical maps are available at NeuroVault.org (https://neurovault.org/collections/MXWQPPCW/). Neuroimaging data necessary to recreate all analyses are available in brain imaging data structure format (BIDS) on OpenNeuro (https://openneuro.org/datasets/ds002994/versions/1.0.1). Behavioural and eye-tracking data, codes for behaviour, eye-tracking and fMRI analysis, and all experiment codes are available on GitHub (https://github.com/GabrielPelletier/fribblesFMRI_object-value-construction).

### Data exclusion

Participants who performed poorly in the bidding fMRI task were excluded from analysis based on preregistered exclusion criteria. Specifically, participants with average rating error ≥ 15 ILS in both conditions, or an average rating error ≥ 15 ILS for any single object were excluded. These criteria ensured that no participant using heuristics to estimate value (i.e. rough guessing based on a reduced number of attributes) was included in the final sample. Eye-tracking data were discarded for a trial if < 70% of samples could be labeled as fixations.

### Statistical analysis

#### Behavioural data analysis

Learning outside the scanner was assessed by the change in average value rating error across learning blocks. Error was defined as the absolute difference between the rating provided by the subject and the true value of the object or attribute. A repeated-measure ANOVA with learning block (5 levels) and condition (2 levels) as within-subject factors was used to analyze error across learning trials. Group-level value rating error in the learning probes was compared between conditions using a paired-sample t-test.

Performance in the bidding task inside the scanner was analyzed by calculating the average error (absolute difference between bid value and instructed value) across the six repetitions for each of the 12 objects, as well as the average error by condition. Group-level bidding error was compared between conditions using a paired-sample t-test. Rating reaction times were similarly compared between conditions.

#### Eye-tracking data analysis

Eye-tracking data files in EyeLink (.edf) format were converted using the Edf2Mat Matlab Toolbox (https://github.com/uzh/edf-converter). Periods of eye blinks were removed from the data, after which the x and y coordinates and the duration of each fixation during the 3 s of object presentation were extracted. We identified each fixation according to whether it fell on one or the other of the learned attributes, or neither. The attribute AOIs were defined by drawing the two largest equal-sized rectangles centered on the attributes of interest that did not overlap with each other. The same two AOIs were used for the 6 objects within each set. All AOIs covered an equal area of the visual field, although the positions varied between object sets. An example of the preregistered AOIs is presented on **Fig. 6*B***. AOIs for all object sets along with their exact coordinates in screen pixels are reported in the preregistration document (osf.io/4d2yr).

For each subject and each condition, we calculated the average number of fixations per trial, and the number of fixations in each of the AOIs. We also calculated the average duration of individual fixations within each AOI and the total time spent fixating on each AOI. Finally, we calculated the average number of transitions from one attribute-AOI to the other. We counted as a transition every instance of a fixation falling on an AOI immediately preceded by a fixation falling on the other AOI. These variables were compared between conditions at the group-level using paired-sample t-tests.

#### fMRI data preprocessing

Raw imaging data in DICOM format were converted to NIfTI format and organized to fit the Brain Imaging Data Structure (BIDS) (Gorgolewski et al., 2016). Facial features were removed from the anatomical T1w images using pydeface (https://github.com/poldracklab/pydeface). Preprocessing was performed using fMRIPprep 1.3.0.post2 ((Esteban et al., 2019), RRID:SCR_016216), based on Nipype 1.1.8 ((Gorgolewski et al., 2011), RRID:SCR_002502).

Anatomical data preprocessing: The T1-weighted (T1w) image was corrected for intensity non-uniformity (INU) with N4BiasFieldCorrection (Tustison et al., 2010), distributed with ANTs 2.2.0 (Avants et al., 2008)(RRID:SCR_004757) and used as T1w-reference throughout the workflow. The T1w-reference was then skull-stripped using antsBrainExtraction.sh (ANTs 2.2.0), using OASIS30ANTs as target template. Brain surfaces were reconstructed using recon-all (FreeSurfer 6.0.1, RRID:SCR_001847, (Dale et al., 1999)), and the brain mask estimated previously was refined with a custom variation of the method to reconcile ANTs-derived and FreeSurfer-derived segmentations of the cortical gray-matter of Mindboggle (RRID:SCR_002438, (Klein et al., 2017)). Spatial normalization to the ICBM 152 Nonlinear Asymmetrical template version 2009c ((Fonov et al., 2009) RRID:SCR_008796) was performed through nonlinear registration with antsRegistration (ANTs 2.2.0), using brain-extracted versions of both T1w volume and template. Brain tissue segmentation of cerebrospinal fluid (CSF), white-matter (WM) and gray-matter (GM) was performed on the brain-extracted T1w using fast (FSL 5.0.9, RRID:SCR_002823, (Zhang et al., 2001)).

Functional data preprocessing: For each of the 8 BOLD runs per subject (across all tasks and sessions), the following preprocessing was performed. First, a reference volume and its skull-stripped version were generated using a custom methodology of fMRIPrep. A deformation field to correct for susceptibility distortions was estimated based on two echo-planar imaging (EPI) references with opposing phase-encoding directions, using 3dQwarp (Cox and Hyde, 1997) (AFNI 20160207). Based on the estimated susceptibility distortion, an unwarped BOLD reference was calculated for a more accurate co-registration with the anatomical reference. The BOLD reference was then co-registered to the T1w reference using bbregister (FreeSurfer) which implements boundary-based registration (Greve and Fischl, 2009). Co-registration was configured with nine degrees of freedom to account for distortions remaining in the BOLD reference. Head-motion parameters with respect to the BOLD reference (transformation matrices, and six corresponding rotation and translation parameters) were estimated before any spatiotemporal filtering using mcflirt (FSL 5.0.9, (Jenkinson et al., 2002)). The BOLD time-series (including slice-timing correction when applied) were resampled onto their original, native space by applying a single composite transform to correct for head-motion and susceptibility distortions. These resampled BOLD time-series will be referred to as preprocessed BOLD in original space, or just preprocessed BOLD. The BOLD time-series were resampled to MNI152NLin2009cAsym standard space, generating a preprocessed BOLD run in MNI152NLin2009cAsym space. First, a reference volume and its skull-stripped version were generated using a custom methodology of fMRIPrep. Several confounding time-series were calculated based on the preprocessed BOLD: framewise displacement (FD), DVARS and three region-wise global signals. FD and DVARS were calculated for each functional run, both using their implementations in Nipype (following the definitions by (Power et al., 2014)). The three global signals were extracted within the CSF, the WM, and the whole-brain masks. Additionally, a set of physiological regressors were extracted to allow for component-based noise correction (CompCor, (Behzadi et al., 2007)). Principal components were estimated after high-pass filtering the preprocessed BOLD time-series (using a discrete cosine filter with 128s cut-off) for the two CompCor variants: temporal (tCompCor) and anatomical (aCompCor). Six tCompCor components are then calculated from the top 5% variable voxels within a mask covering the subcortical regions. This subcortical mask is obtained by heavily eroding the brain mask, which ensures it does not include cortical GM regions. For aCompCor, six components are calculated within the intersection of the aforementioned mask and the union of CSF and WM masks calculated in T1w space, after their projection to the native space of each functional run (using the inverse BOLD-to-T1w transformation). The head-motion estimates calculated in the correction step were also placed within the corresponding confounds file. All resamplings were performed with a single interpolation step by composing all the pertinent transformations (i.e. head-motion transform matrices, susceptibility distortion correction, and co-registrations to anatomical and template spaces). Gridded (volumetric) resamplings were performed using antsApplyTransforms (ANTs), configured with Lanczos interpolation to minimize the smoothing effects of other kernels (Lanczos, 1964). Non-gridded (surface) resamplings were performed using mri_vol2surf (FreeSurfer).

Confound files were created for each scan (each run of each task of each participant, in .tsv format), with the following columns: standard deviation of the root mean squared (RMS) intensity difference from one volume to the next (DVARS), six anatomical component based noise correction method (aCompCor), frame-wise displacement, and six motion parameters (translation and rotation each in 3 directions) as well as their squared and temporal derivatives (Friston 24-parameter model (Friston et al., 1996)). A single time point regressor (a single additional column) was added for each volume with FD value larger than 0.9, in order to model out volumes with excessive motion. Scans with more than 15% scrubbed volumes were excluded from analysis.

#### fMRI data analysis

fMRI data were analyzed using FSL FEAT (fMRI Expert Analysis Tool) of FSL (Smith et al., 2004). A general linear model (GLM) was estimated to extract contrasts of parameter estimate at each voxel for each subject for each of the four fMRI runs (first level analysis). Contrasts of parameter estimate from the four runs were then averaged within participants using a fixed effect model (second level analysis). Group-level effects were estimated using a mixed effect model (FSL’s FLAME-1).

General linear model: The GLM included one regressor modelling the 3-s object presentation time for configural trials, and one regressor modelling object presentation for elemental trials. The model also included one regressor modelling object presentation for the configural trials modulated by the value rating of the object provided on each trial (mean centered), and the equivalent regressor for elemental trials. We included four regressors modelling the rating epoch of the trial, with two unmodulated regressors modelling the rating scale for configural trials and elemental trials separately, and two regressors modelling the rating scale epoch modulated by value ratings (mean-centered) for configural trials and elemental trials separately. The duration of the rating event in these four regressors was set to the average rating reaction time across all participants and runs. Rating reaction times were accounted for in the model using a separate regressor modelling the rating epoch for all trials, modulated by the trial-wise reaction time (mean-centered). The duration was set to the maximum response time of 3 s in cases where the time limit was reached. All regressors included in this GLM were convolved with a canonical double-gamma hemodynamic response function. Their temporal derivatives were also included in the model, with the motion and physiological confounds estimated by fMRIPrep as described above.

#### Regions of interest (ROI)

A vmPFC ROI was defined using the combination of the Harvard-Oxford regions frontal pole, frontal medial cortex, paracingulate gyrus and subcallosal cortex, falling between MNI x = −14 and 14 and z < 0, as in (Schonberg et al., 2014). This ROI was used for small volume correction where specified.

In addition, we defined four ROIs along the ventral visual stream of the brain; the perirhinal cortex (PRC), parahippocampal place area (PPA), fusiform face area (FFA) and the lateral occipital complex (LOC) using functional localizer data, as in (Erez et al., 2016). The PRC was defined based on a probabilistic map (Devlin and Price, 2007) created by superimposing the PRC masks of 12 subjects, segmented based on anatomical guidelines in MNI-152 standard space. We thresholded the probabilistic map to keep voxels having more than 30% chance of belonging to the PRC, as in previous work (Erez et al., 2016). The lateral occipital complex (LOC) was defined as the region located along the lateral extent of the occipital pole that responded more strongly to objects than scrambled objects (p < 0.001, uncorrected). The fusiform face area (FFA) was defined as the region that responded more strongly to faces than objects. The PPA was defined as the region that responded more strongly to scenes than to objects. For each of these contrasts, a 10 mm radius sphere was drawn around the peak voxel in each hemisphere using FSL (fslmaths). To analyze brain activity in these regions during the bidding task, cope images from the second-level analysis (average of the four runs for each participant) were converted to percent signal change (as described in (Mumford, 2007)), before averaging across all voxels within each ventral visual stream ROI. Group-level activations were compared against 0 using one-sample t-tests.

#### Functional connectivity analysis

Functional connectivity was assessed using generalized psychophysiological interaction analysis (gPPI) to reveal brain regions where BOLD time-series correlate significantly with the time-series of a target seed region in one condition more than another (McLaren et al., 2012). The seed region was defined based on the significant activation cluster found in the group-level analysis for the configural trials value-modulation contrast, small-volume corrected for the vmPFC ROI (**Fig. 4*A***). The seeds’ neural response to configural and elemental trials were estimated by deconvolving the mean BOLD signal of all voxels inside the seed region (Gitelman et al., 2003).

The gPPI-GLM included the same regressors as the main GLM described above, plus two psychophysiological interaction (PPI) regressors of interest: one regressor modelling the seed region’s response to configural trials, and one regressor modelling the seed region’s response to elemental trials. These regressors were obtained by multiplying the seed region time-series with an indicator function for object presentation of the corresponding condition, and then re-convolving the result with the double-gamma hemodynamic function. The model additionally included one regressor modelling the BOLD time-series of the seed region.

### Inference criteria

For behavioural and eye-tracking analysis, we used the standard threshold of *p* < 0.05 for statistical significance, and we report exact *p*-values and effect sizes for all analyses. Neuroimaging data are reported at the group level with statistical maps thresholded at *Z* > 3.1 and cluster-based Gaussian Random Field corrected for multiple comparisons with a (whole brain corrected) cluster significance threshold of *p* < 0.05. We report analyses restricted to the vmPFC ROI using the same inference criteria, with increased sensitivity to detect effects in this region defined *a priori* due to fewer comparisons (small volume correction). Ventral visual stream ROI results are reported using the statistical threshold of *p* < 0.05, Bonferroni-corrected for four comparisons (the number of ROIs) (*p* < 0.0125).

### Deviations from preregistration

The most substantial deviation from the preregistered analysis concerns the main GLM defined for fMRI analysis. We controlled for reaction times differently than what was stated in the preregistration, this was done due to a mistake in the preregistered analysis plan that proposed an approach different from the usual process of accounting for RT (Schonberg et al., 2014; Botvinik-Nezer et al., 2020; Salomon et al., 2020). We also carried out supplementary fMRI analyses including accuracy confound regressors in the GLM after behavioural analysis revealed a trend difference in accuracy between conditions. This analysis did not yield substantially different results, and we thus report results from the model without accuracy regressors, as preregistered.

## RESULTS

### Behaviour

We first present the behavioural results from the behavioural and fMRI studies, to establish the replicability of the behavioural effects.

#### Learning phase

Participants learned the value of novel multi-attribute objects under two conditions, elemental and configural. Learning behaviour differed between conditions in both the behavioural and the MRI sample (this phase of the task was performed outside the scanner in both studies), with configural associations being generally harder to learn than elemental ones, as detailed below.

Value rating errors decreased across learning blocks and were overall higher in the configural condition (**Fig. 3*A***). A repeated measures ANOVA with block and condition as within-subject factors, revealed a main effect of block (behavioural sample F_4, 128_ = 58.21, *p* < 0.001, η^2^_p_ = 0.45; fMRI sample F_4, 164_ = 60.73, *p* < 0.001, η^2^_p_ = 0.40) and a main effect of condition (behavioural sample F_1, 32_ = 372.14, *p* < 0.001, η^2^_p_ = 0.56; fMRI sample F_1, 41_ = 470.84, *p* < 0.001, η^2^_p_ = 0.56) on value rating error. We also found a significant block by condition interaction (behavioural sample F_4, 128_ = 37.98, *p* < 0.001, η^2^_p_ = 0.35; fMRI sample F_4, 164_ = 30.20, *p* < 0.001, η^2^_p_ = 0.25). This interaction reflects that error rates were more similar across conditions as learning wore on, although rating error remained significantly greater in the configural compared to the elemental condition on the last (fifth) learning block (paired-sample T-test, behavioural sample t_32_ = 4.69, *p* < 0.001, Cohen’s *d* = 0.817; fMRI sample t_41_ = 6.46, *p* < 0.001, Cohen’s *d* = 0.90).

**Figure 3.**
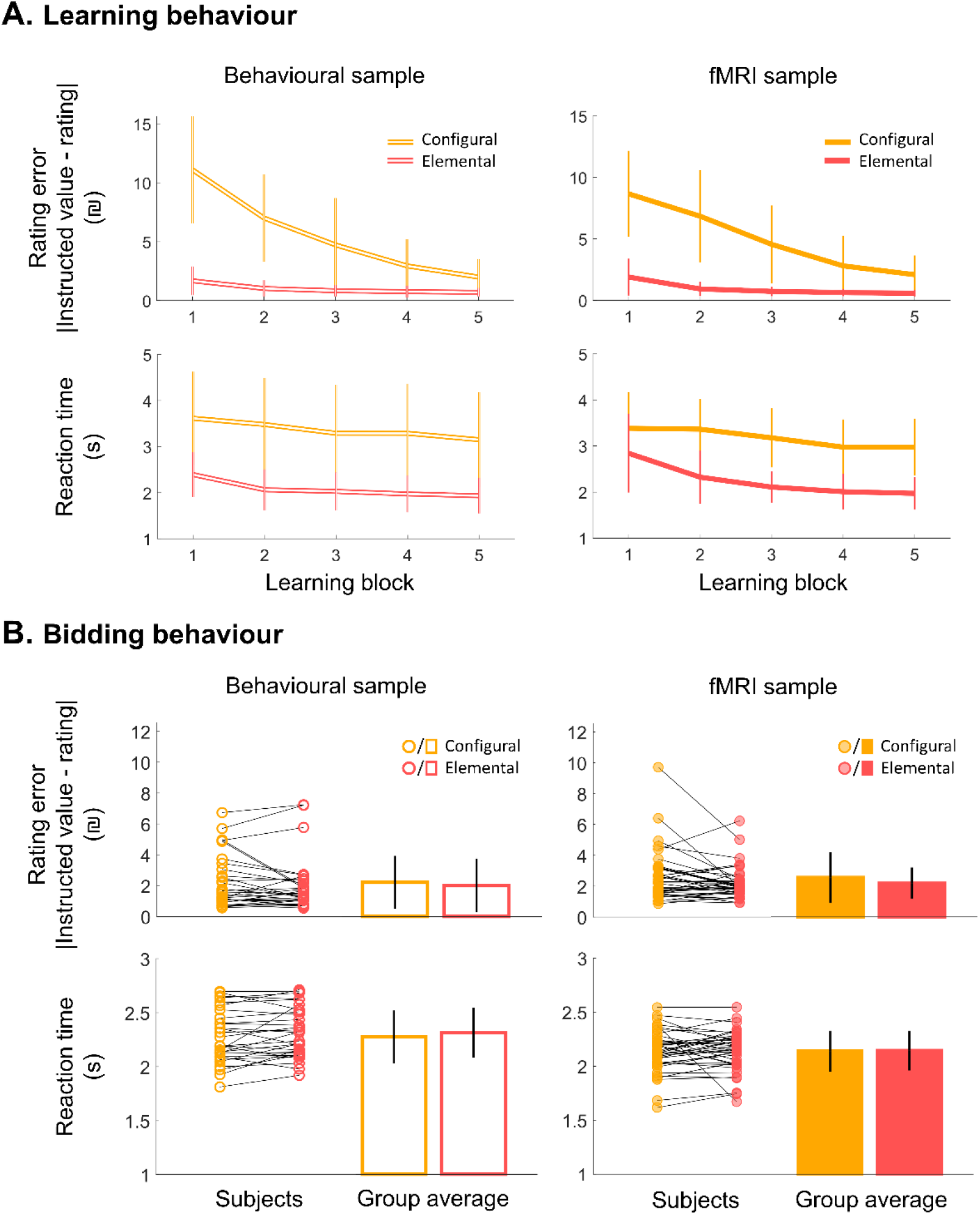
Behaviour in the learning and bidding phases. **A)** Accuracy (top) and reaction time (bottom) across learning blocks in configural and elemental conditions. **B)** Individual and group average value rating error (top) and reaction time (middle) in the fMRI bidding phase collapsed across all trials for each condition. Accuracy is measured in terms of rating error, corresponding to the absolute difference between value rating and the instructed value of the fribble, averaged across all trials within a learning block or within the bidding phase, by condition. Instructed value corresponds to the value of the single salient attribute in the learning blocks in the elemental condition, and to the sum of the two attributes in the bidding phase. In the learning blocks and bidding phase in the configural condition, instructed value corresponds to the value associated with the configuration of two attributes. Error bars represent one standard deviation from the group mean.

Reaction times also decreased across learning blocks (main effect of block, behavioural sample F_4, 128_ = 7.17 *p* < 0.001, η^2^_p_ = 0.09; fMRI sample F_4, 164_ = 26.38, *p* < 0.001, η^2^_p_ = 0.22). Reaction times were significantly faster in the elemental compared to the configural condition (main effect of condition, behavioural sample F_1, 32_ = 467.58, *p* < 0.001, η^2^_p_ = 0.62; fMRI sample F_1, 41_ = 391.35, *p* < 0.001, η^2^_p_ = 0.51). There was no significant block by condition interaction in the behavioural sample (F_4, 128_ = 0.387, *p* = 0.818, η^2^_p_ = 0.005), but there was a significant interaction in the fMRI sample (F_4, 164_ = 4.35, *p* = 0.002, η^2^_p_ = 0.05).

After five learning blocks, participants completed a learning probe without feedback outside the scanner. The learning probe was designed to assess the ability to assign value to the objects during extinction. It was also important to assess the ability to sum two attribute values in the elemental condition, which only included single attribute-value associations in the learning blocks. In the learning probe, accuracy was lower in the elemental condition compared to the configural condition in the behavioural sample (paired-sample T-test; t_32_ = 2.13, *p* = 0.041, Cohen’s *d* = 0.372) but was not significantly different between conditions in the fMRI sample (t41 = 1.30, *p* = 0.201, Cohen’s *d* = 0.201). Participants were slower in the elemental compared to the configural condition in both samples (behavioural sample t_32_ = 5.47, *p* < 0.001, Cohen’s *d* = 0.953; fMRI sample t_41_ = 9.56, *p* < 0.001, Cohen’s *d* = 1.48).

#### Bidding task

After learning, participants were shown objects from the configural and elemental sets and were asked to bid. Participants in the fMRI study performed the learning phase outside the scanner and then performed the bidding stage while scanned with fMRI. Bidding accuracy was high and not significantly different between the configural (mean rating error = 2.26NIS, SD = 1.66) and elemental (M = 2.04NIS, SD = 1.66) conditions for the behavioural sample (t_32_ = 1.08, *p* = 0.289, Cohen’s *d* = 0.188) (**Fig. 3*B***). In the fMRI sample, bids tended to be closer to the instructed value (smaller error) in the elemental (M = 2.18NIS, SD = 1.02) compared to the configural condition (M = 2.55NIS, SD = 1.63), although the difference did not reach significance and the effect was marginal (t41 = 1.90, *p* = 0.065, Cohen’s *d* = 0.293). Value rating reaction times were not significantly different between conditions (behavioural sample t_32_ = 1.80, *p* = 0.081, Cohen’s *d* = 0.314; fMRI sample t_41_ = 0.251, *p* = 0.803, Cohen’s *d* = 0.038). Thus, despite some behavioural differences between conditions in the learning phase, accuracy and reaction times were similar across conditions in the bidding phase, which was the focus of subsequent analyses.

### fMRI signal in the vmPFC selectively tracks configural object value

We hypothesized that the fMRI signal in vmPFC would correlate with configural object value, and that the correlation of vmPFC signal and value would be stronger for configural compared to elemental trials. To test this hypothesis, we preregistered analysis of value modulation effects at the time of object presentation in the *a priori* defined vmPFC region of interest using small volume correction. The hypothesized value signal in the vmPFC was not detected during the object presentation epoch but was instead evident at the time of value rating. Two clusters in the vmPFC were significantly correlated with value for configural trials in the rating phase (**Fig. 4*A***). In contrast, no activation clusters were found to correlate with value in the elemental trials, and the direct condition contrast revealed a significant condition by value interaction in the vmPFC, in which signal was correlated more strongly with value in configural compared to elemental trials (**Fig. 4*B***).

**Figure 4.**
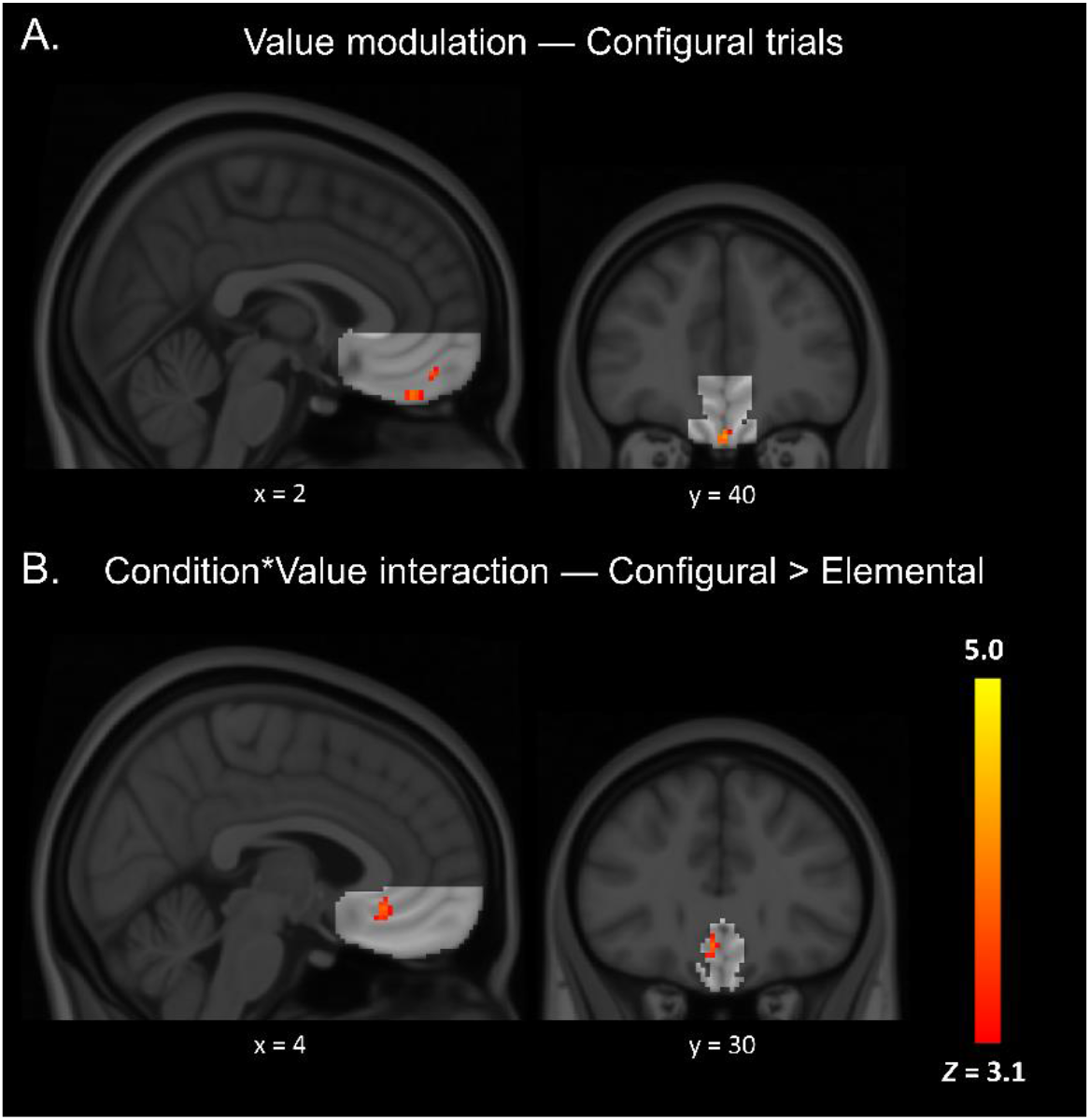
Value-modulated activation clusters during value rating in the vmPFC. **A)** Clusters where the fMRI signal was significantly modulated by value in configural trials. **B)** Cluster where value modulation was stronger for configural compared to elemental trials. Results were small volume corrected (SVC) for the preregistered vmPFC region of interest (bright area), at a cluster-forming threshold of *Z* > 3.1 and *p* < 0.05. The color bar indicates Z-statistics. Numbers below slices indicate MNI coordinates.

### Condition by value interaction in the ventral visual stream

We next tested whether the ventral visual stream ROIs were sensitive to the valuation condition. Our preregistered hypothesis was that at the time of object presentation, fMRI signals in the PRC, and not in anterior VVS regions, would be greater in response to objects learned in the configural condition. We found no significant main effect of condition on BOLD in the PRC (*p* = 0.460) or any other VVS region (LOC *p* = 0.286; FFA *p* = 0.731; PPA *p* = 0.136) (**Fig. 5*B***) at the time of object presentation, indicating that during this time, VVS ROIs were similarly activated in response to objects learned in the configural and elemental conditions.

**Figure 5.**
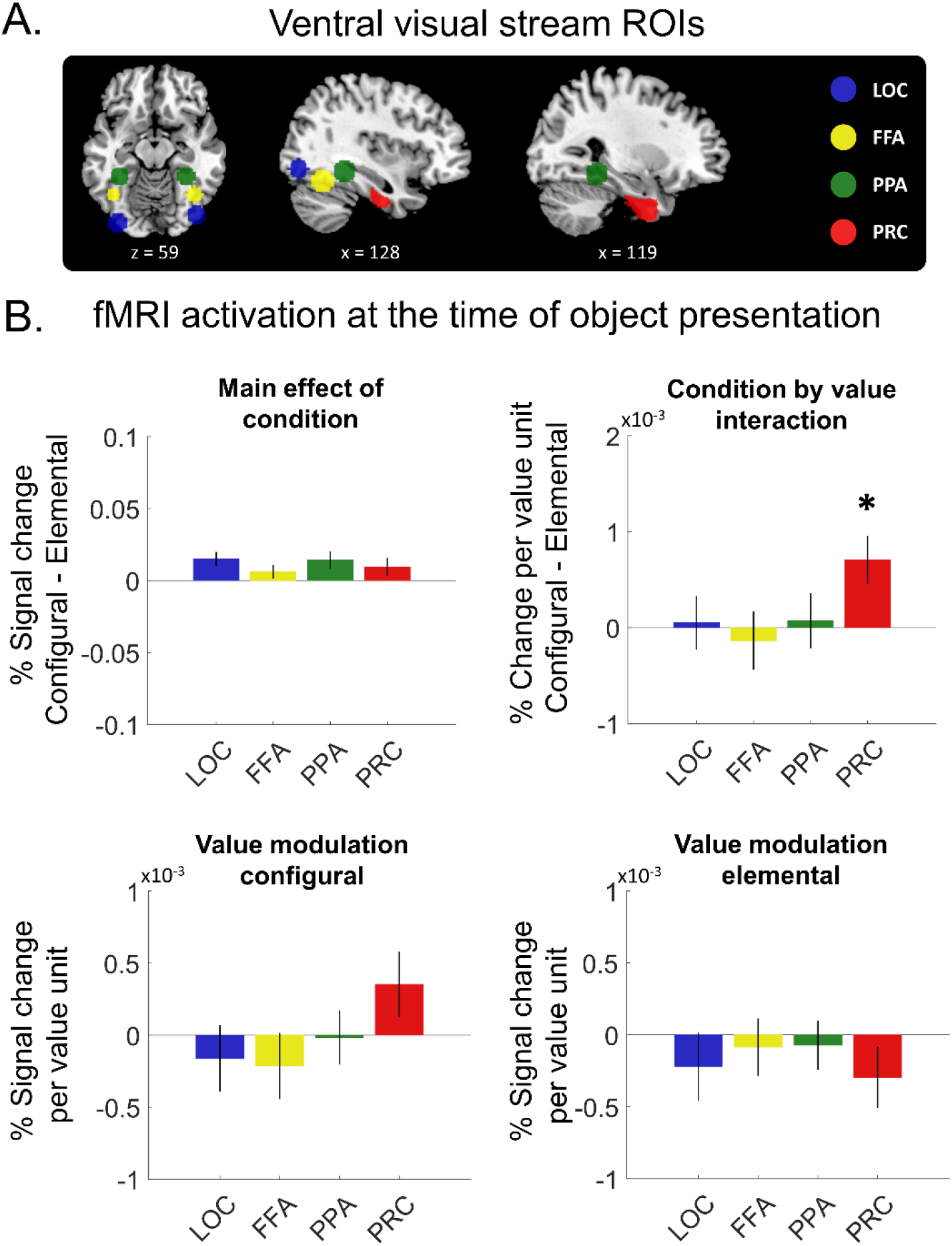
Ventral visual stream regions of interest analysis. **A)** Regions of interest. The lateral occipital complex (LOC), fusiform face area (FFA) and parahippocampal place area (PPA) ROIs shown for a representative participant. The perirhinal cortex (PRC) ROI was the same for all participants. Numbers indicate coordinates in MNI space. **B)** Percent signal change during the object presentation epoch. The top-left panel shows the main effect of condition, assessed with the configural minus elemental trials contrast. The top-right panel shows the condition by value interaction, assessed by contrasting the effect of value modulation in configural trials, minus the effect of value modulation in elemental trials. Bottom panels show the value modulation effect for configural and elemental trials separately. Error bars represent SEM. Asterisk indicates significance at *p* < 0.05 for one sample t-test against 0, after Bonferroni correction for four comparisons.

We next examined whether VVS regions were sensitive to value. We found a significant condition by value interaction in the PRC: in this region, the BOLD signal associated with value was stronger for configural compared to elemental trials (*p* = 0.016, Bonferroni corrected for four ROIs) (**Fig. 5*B***). This effect was specific to the PRC and was not found in more posterior regions of the VVS (LOC, FFA and PPA uncorrected *ps* > 0.727). We decomposed the interaction by examining value modulation in configural and elemental trials separately. In the PRC, there was a non-significant trend in BOLD signal to be positively correlated with value in configural trials (uncorrected *p* = 0.127), and negatively correlated with value in elemental trials (uncorrected *p* = 0.172) (**Fig. 5B**). There was no significant effect of condition (uncorrected *ps* > 0.216) and no condition by value interaction (uncorrected *ps* > 0.394) in any VVS regions during the value rating epoch.

### Whole-brain examination of configural and elemental evaluation

ROI analyses revealed value modulation effects in the vmPFC and a condition by value interaction in the PRC, but no main effect of condition. We next carried out whole-brain analyses to ask which brain regions, if any, were on average more active in configural in contrast to elemental trials regardless of value at the time of object presentation, shown in **Table 1**. **Table 2** shows the clusters for the opposite contrast.

**Table 1.** Clusters with significantly more activity during configural compared to elemental object presentation

| Cluster | Voxels | Anatomic region included in cluster | Voxels in region | Peak Z | MNI (x, y, z) | P |
| --- | --- | --- | --- | --- | --- | --- |
| 1 | 67 | Middle Frontal Gyrus (R) | 55 | 4.53 | 46, 14, 50 | 0.014 |
| 2 | 78 | Caudate (R) | 34 | 4.88 | 8, 6, 6 | 0.006 |
|  |  | Thalamus (R) | 11 |  |  |  |
| 3 | 171 | Precuneus Cortex (R) | 160 | 4.17 | 14, -68, 28 | 1.06e-05 |
| 4 | 206 | Inferior Frontal Gyrus pars triangularis (R) | 54 | 4.84 | 56, 28, 22 | 1.37e-06 |
|  |  | Middle Frontal Gyrus (R) | 45 |  |  |  |
|  |  | Inferior Frontal Gyrus pars opercularis (R) | 13 |  |  |  |
|  |  | Frontal Pole (R) | 9 |  |  |  |
| 5 | 259 | Superior Lateral Occipital Cortex (L) | 173 | 4.19 | -40, -70, 44 | 5.96e-08 |
|  |  | Angular Gyrus (L) | 38 |  |  |  |
| 6 | 348 | Superior Lateral Occipital Cortex (R) | 282 | 4.64 | 42, -74, 48 | 8.64e-10 |
|  |  | Angular Gyrus (R) | 14 |  |  |  |
| 7 | 426 | Superior Frontal Gyrus (L) | 207 | 5.42 | -2, 36, 42 | 2.36e-11 |
|  |  | Paracingulate Gyrus (L) | 120 |  |  |  |
|  |  | Superior Frontal Gyrus (R) | 32 |  |  |  |
|  |  | Paracingulate Gyrus (R) | 21 |  |  |  |
| 8 | 429 | Cerebellum (R) | 361 | 5.34 | -14, -82, -34 | 2.06e-11 |
| 9 | 623 | Cerebellum (L) | 511 | 5.76 | 14, -72, -30 | 6.27e-15 |
| 10 | 1081 | Precuneous Cortex (L) | 673 | 4.99 | 2, -68, 60 | 5.07e-22 |
|  |  | Precuneous Cortex (R) | 154 |  |  |  |
|  |  | Cuneal Cortex (L) | 48 |  |  |  |
|  |  | Cuneal Cortex (R) | 9 |  |  |  |
|  |  | Supracalcarine Cortex (L) | 7 |  |  |  |
| 11 | 1381 | Frontal Pole (L) | 471 | 5.11 | -38, 22, -4 | 4.11e-26 |
|  |  | Middle Frontal Gyrus (L) | 195 |  |  |  |
|  |  | Inferior Frontal Gyrus pars triangularis (L) | 145 |  |  |  |
|  |  | Frontal Orbital Cortex (L) | 136 |  |  |  |
|  |  | Insular Cortex (L) | 32 |  |  |  |
|  |  | Inferior Frontal Gyrus pars opercularis (L) | 16 |  |  |  |

Results were whole-brain cluster-corrected, at a cluster-forming threshold of $Z > 3.1$ and $p < 0.05$ .

**Table 2.** Clusters with significantly more activity during elemental compared to configural object presentation

| Cluster | Voxels | Anatomic region included in cluster | Voxels in region | MNI (x, y, z) | Peak Z | p |
| --- | --- | --- | --- | --- | --- | --- |
| 1 | 76 | Precentral Gyrus (R) | 76 | 54, -2, 40 | 5.29 | 0.007 |
| 2 | 207 | Supplementary Motor Cortex (L) | 164 | -6, 2, 64 | 6.08 | 1.31e-06 |
|  |  | Superior Frontal Gyrus (L) | 17 |  |  |  |
| 3 | 308 | Precentral Gyrus (L) | 206 | -64, 8, 22 | 5.39 | 6.08e-09 |
|  |  | Inferior Frontal Gyrus pars opercularis (L) | 19 |  |  |  |
| 4 | 485 | Precentral Gyrus (L) | 362 | -56, -4, 38 | 5.57 | 1.79e-12 |
|  |  | Postcentral Gyrus (L) | 15 |  |  |  |

Results were whole-brain cluster-corrected, at a cluster-forming threshold of $Z > 3.1$ and $p < 0.05$ .

### Eye movements distinguish between configural and elemental evaluation

In the previous section, we report different brain regions recruited in configural and elemental object evaluation. We next investigated whether eye movements during the same 3-s object presentation epoch of the bidding task trials were different between conditions (**Fig. 6*A*, Fig. 6*B***). The average number of fixations made on the whole object was similar across conditions (behavioural sample t_32_ = 1.741, *p* = 0.091, Cohen’s *d* = 0.303; fMRI sample t_32_ = 0.479, *p* = 0.635, Cohen’s *d* = 0.083). However, we found consistent condition differences across the two samples in eye movements with respect to fixations to the value-predictive attributes. Participants made significantly more transitions between these attributes in the configural compared to the elemental condition (behavioural sample t_32_= 3.364, *p* = 0.002, Cohen’s *d* = 0.586; fMRI sample t_32_ = 2.659, *p* = 0.012, Cohen’s *d* = 0.463), and the average duration of individual fixations was longer in the elemental condition (behavioural sample t_32_ = 3.611, *p* = 0.001, Cohen’s *d* = 0.559; fMRI sample t_32_ = 2.211, *p* = 0.034, Cohen’s *d* = 0.385).

Given this observation, we carried out exploratory (not preregistered) analyses to investigate whether gaze differences between conditions were related to differences in the brain. For each participant, we calculated the percent signal change for the configural minus elemental contrast, averaged across voxels of all significant clusters more active in configural compared to elemental trials from the group level whole brain analysis (**Table 1**). We correlated this brain activation variable with the difference between the average number of gaze transitions made in the configural minus the elemental trials. This revealed a significant positive correlation between the two variables: the greater the difference in brain activations, the greater the difference in eye movements (Pearson r = 0.425, *p* = 0.015) (**Fig. 6C**). There was no significant correlation between brain activation and the difference in average durations of fixations (Pearson r = −0.075, *p* = 0.682) or the difference between the number of fixations (Pearson r = 0.110, *p* = 0.550).

**Figure 6.**
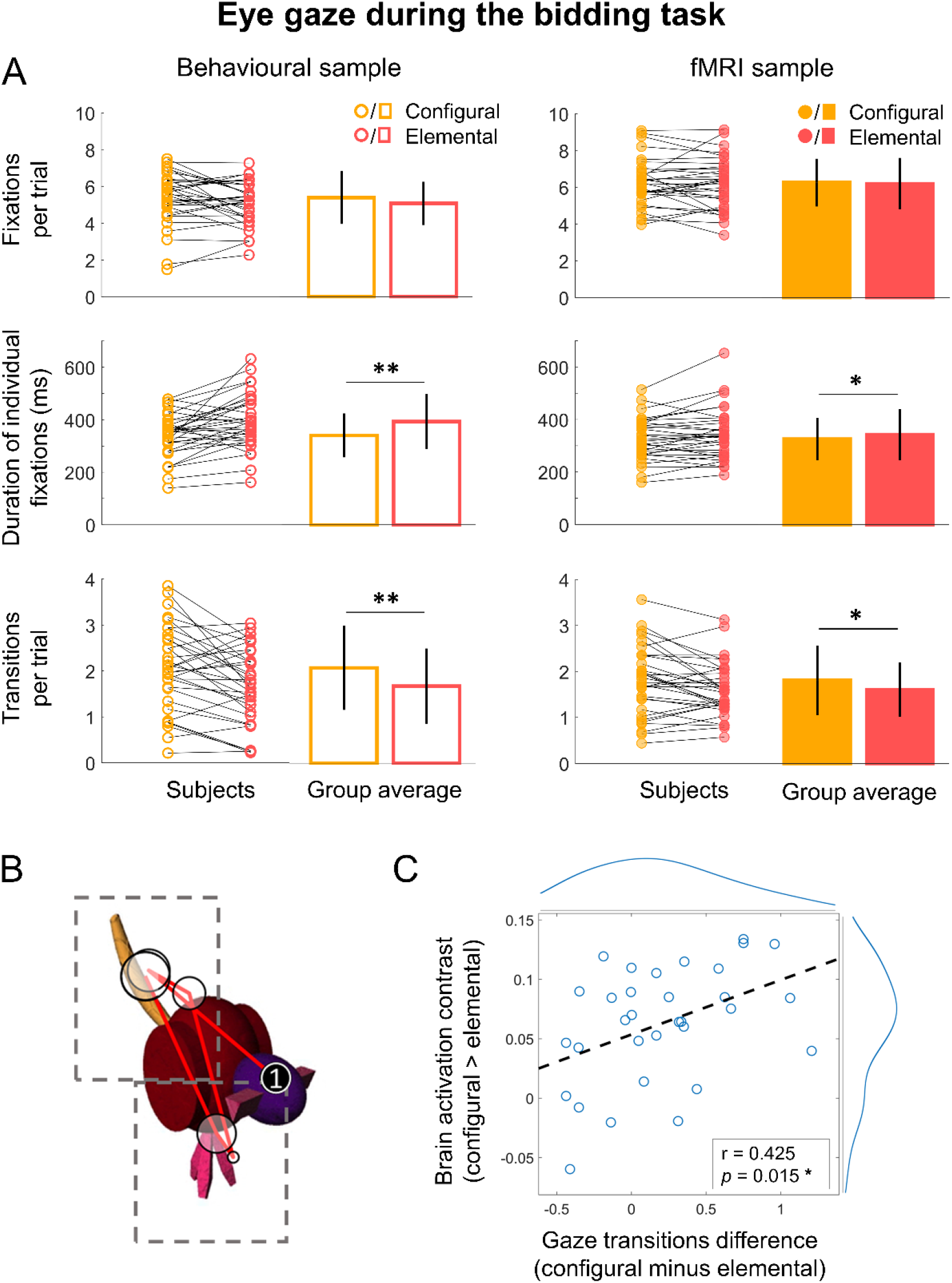
Eye-tracking results. **A)** Average number of fixations per trial (top), average duration of individual fixations on attribute-AOIs (middle) and average number of transitions between attribute-AOIs per trial (bottom). Error bars represent one standard deviation from the group mean. * indicate significant differences between conditions at *p* < 0.05, ** *p* < 0.01. **B)** Example of eye gaze data for one representative trial. Circles represent fixations, and their diameter indicate the relative fixation duration. The first fixation is identified (1) with the subsequent fixations of the scan path indicated by the red lines. Dashed boxes represent areas of interest for the two value-relevant attributes. In this example, there were six fixations and two transitions (the scan path crosses twice from one attribute AOI to the other). **C)** Significant correlation between gaze transitions and brain activations across participants of the fMRI sample. The y-axis displays the participant-level activation contrast for configural minus elemental trials at the time of object presentation, averaged across the brain voxels displaying a significant main effect of condition at the group-level (**Table 1**). Data points represent individual participants, superimposed with the best-fitting linear regression line (dashed); ‘r’ value indicates Pearson correlation. Curves on the top and right hand-side of the plot represent the distribution of the x- and y-axis variables, respectively.

### Functional connectivity analysis

We carried out a preregistered functional connectivity analysis using gPPI, defining the seed as the significant vmPFC clusters found for configural trials value modulation (**Fig. 4*A***). The gPPI analysis did not reveal any clusters across the whole-brain, and no VVS region displaying evidence of greater functional connectivity with the vmPFC seed in configural compared to elemental trials, or vice versa.

## DISCUSSION

Here, using both eye-tracking and fMRI, we show behavioral and neural evidence for two distinct mechanisms of assessing the value of multi-attribute objects. We found that evaluation of complex objects relied on different patterns of information acquisition, indexed by eye movements, and engaged different brain regions when value was predicted by configural relationships between attributes compared to when value could be summed from the independent values of individual attributes. Activity in the perirhinal cortex was correlated with value in configural more than elemental trials during object presentation, whereas at the time of value rating, vmPFC showed value-modulated signal for configural trials only. Participants made longer fixations on individual attributes in the elemental condition and made more gaze transitions from one attribute to another when viewing objects in the configural condition. Moreover, at the participant level, the between-condition difference in the number of gaze transitions was correlated with the difference in brain activation.

These experiments in three different samples provide evidence converging with the findings from a recent study in patients with vmPFC damage using the same stimuli (Pelletier and Fellows, 2019). That lesion study found that vmPFC damage impaired binary decisions between fribbles in the configural condition, but not in the elemental condition. The current work provides additional support for the hypothesis that vmPFC has a unique role in inferring the value of objects based on configural information: BOLD signal in that region was only detectably modulated by object value in the configural and not the elemental condition. The present study further argues that evaluation in the configural condition engages the PRC, a region known to be critical for multi-attribute object recognition, but here for the first time also implicated in the evaluation of such objects.

We did not find that the total value of an object obtained by combining two separately learned attribute-values was reflected in the vmPFC fMRI signal. This null result alone cannot rule out that vmPFC is involved in value integration from multiple elements. However, taken together with the finding that damage to vmPFC did not substantially impair the ability to make choices based on such values, it suggests the existence of alternate mechanisms for value construction under such conditions, not requiring vmPFC.

Across published fMRI work, vmPFC is reliably associated with subjective value (Rushworth and Behrens, 2008; Bartra et al., 2013). Activity in the vmPFC has also been shown to reflect the values of items composed of multiple attributes, each modelled as independently predictive of value (Basten et al., 2010; Lim et al., 2013; Suzuki et al., 2017). Although the assumption of elemental value integration made in these studies was consistent with their data, it is possible that whole option-values were nonetheless estimated ‘configurally’. No previous work has contrasted these distinct types of multi-attribute valuation, leaving unclear whether vmPFC value signals reflect elemental or configural assessment. The current findings add to the view that the vmPFC is not critical for value integration in general, but rather becomes necessary under a narrower set of conditions (Vaidya and Fellows, 2020). We propose a more specific account whereby the vmPFC is required for inferring value from the configural relationships among lower-level attributes. This view might explain prior observations that patients with vmPFC damage are able to evaluate complex social or aesthetic stimuli, but seem to draw on different information to assess the value of such stimuli, compared to healthy participants (Xia et al., 2015; Vaidya et al., 2018).

The current work also addressed whether regions known to be involved in complex object recognition are likewise involved in assessing the values of such options. We found that fMRI VVS signals were differently sensitive to value across conditions. Specifically, activity in the PRC was modulated by value more for configural compared to elemental trials. There are previous reports of value-correlated signal across the VVS, including in the primary visual cortex (Serences, 2008; Nelissen et al., 2012), lateral occipital complex (Persichetti et al., 2015), the PRC (Mogami and Tanaka, 2006) and several of these regions combined (Arsenault et al., 2013; Kaskan et al., 2017). Across studies, reward has been paired with stimuli ranging in complexity from simple colored gratings to complex objects, but no work previously contrasted conditions in which evaluation relied on characteristics represented at different stages of the VVS hierarchy. Our findings suggest a selective involvement of the PRC in encoding value when it is associated with the high-level (i.e. configural) object representations that this region supports. This is compatible with previous findings that the change in value of faces and objects is associated with changes in face and object processing regions, respectively (Botvinik-Nezer et al., 2020; Salomon et al., 2020), arguing that value learning and storage occurs partly through experience-dependent plasticity in sensory cortex (Schonberg and Katz, 2020).

A condition by value interaction for PRC activity was only observed during object presentation, on average six seconds before value rating, when value-related signals were detected in the vmPFC, arguing against the possibility that value-related PRC activation is driven by the vmPFC. This finding rather suggests that the VVS, in addition to being involved in value learning, is also involved in developing value representations during object recognition, with vmPFC activation following later in the decision process. This is consistent with electrophysiological recordings in macaques, which reported value sensitivity in PRC neurons at about 200 ms after stimulus onset (Mogami and Tanaka, 2006), whereas other work detected value selective signals only after 400-500 ms in the OFC (Wallis and Miller, 2003; Kennerley et al., 2008). Electroencephalography recordings in humans likewise revealed value-correlated signals in response to reward-paired objects emerging earlier in the occipital cortex than in the prefrontal cortex (Larsen and O’Doherty, 2014). With these data, the current work is compatible with the idea that value representations emerge gradually from the onset of sensory processing (Yoo and Hayden, 2018).

We also found systematic differences in eye gaze patterns between conditions, replicated in two samples. Moreover, we found that the greater the difference in gaze transitions in the configural compared to the elemental trials, the greater the difference in brain activation between conditions. Sequential sampling models have shown that value and gaze interact in driving the decision process (Krajbich et al., 2010), and that gaze duration has a causal influence on value (Shimojo et al., 2003). However, little is known about fixation patterns within multi-attribute objects during choice (Krajbich, 2019), and how they relate to the value construction process. Consumer research has extensively studied value construction strategies using process tracing measures including eye-tracking (Russo and Dosher, 1983; Bettman et al., 1998). However, these studies decomposed options by laying out attributes as text and numbers in a table format, which might not relate to the mechanisms underlying everyday choices between complex objects that are likely to be more readily represented in VVS. Here, we provide evidence that when evaluating complex objects having well-controlled visual properties, equal numbers of value-informative attributes, and the same overall value, value construction *per se* is reflected in eye movements and brain activations. These distinct forms of multi-attribute evaluation may inform further work to fully understand the interplay between gaze patterns and value construction during complex decision-making (Busemeyer et al., 2019).

We did not find evidence for increased functional connectivity between the vmPFC and PRC during configural object valuation. This null result must be interpreted with caution, as the study was not powered to find such an effect. There are anatomical connections (Heide et al., 2013) and there is evidence of functional connectivity (Andrews-Hanna et al., 2014) between the vmPFC and the medial temporal lobe in humans. The PRC and medial OFC are reciprocally connected in macaques (Kondo et al., 2005), and disconnecting these regions disrupts value estimation of complex visual stimuli (Clark et al., 2013; Eldridge et al., 2016). The current findings of value-related activations at different stages of the trial in the PRC and the vmPFC suggest that interactions between these two regions might be important for value estimation in configural conditions.

Although we attempted to match the two conditions for difficulty, and further addressed this potential confound by controlling for trial-by-trial rating reaction times and accuracy in fMRI analyses, one limitation of this study is that we could not account for potential condition differences in speed of evaluation during the fixed object presentation time and the subsequent ITI. The slider response requirement also meant that motor responses were confounded with rated values, potentially explaining why motor regions showed value-correlated activation at the time of object presentation.

In conclusion, this neuroimaging study, directly linked to our recent work in lesion patients, provides evidence for two ways of building the value of complex objects, supported by at least partly distinct neural mechanisms. Leveraging object-recognition research to inform studies of multi-attribute value-based decisions, this work suggests that the relationship between attributes and value might influence how an object is processed through the VVS. Research at the interface of these two fields of research may bring novel perspectives on a unified brain basis of both perception and motivated behavior.

## Acknowledgements

This work was supported by grants from the Israel Science Foundation (1798/15 and 2004/15) to Tom Schonberg and from the Canadian Institutes of Health Research (MOP-11920) and the Natural Sciences and Engineering Research Council of Canada (RGPIN-2016-06066) to Lesley Fellows. Gabriel Pelletier was supported by the Zavalkoff Family Foundation as part of the Brain@McGill and Tel Aviv University collaboration, and by a “Sandwich” Scholarship from the Council for Higher Education in Israel. The authors would like to thank Tom Salomon and Shiran Oren for helpful discussions on study design and data analysis, and Anastasia Saliy Grigoryan for help with participant recruitment. We would also like to thank the staff of The Alfredo Federico Strauss Center for Computational Neuroimaging of Tel Aviv University for help with data collection.

## Notes

### Competing Interest Statement

The authors have declared no competing interest.

## REFERENCES

Andrews-Hanna JR, Smallwood J, Spreng RN (2014) The default network and self-generated thought: component processes, dynamic control, and clinical relevance. Ann N Y Acad Sci 1316:29–52.

Arsenault JT, Nelissen K, Jarraya B, Vanduffel W (2013) Dopaminergic Reward Signals Selectively Decrease fMRI Activity in Primate Visual Cortex. Neuron 77:1174–1186.

Avants BB, Epstein CL, Grossman M, Gee JC (2008) Symmetric Diffeomorphic Image Registration with Cross-Correlation: Evaluating Automated Labeling of Elderly and Neurodegenerative Brain. Med Image Anal 12:26–41.

Bartko SJ, Winters BD, Cowell RA, Saksida LM, Bussey TJ (2007) Perirhinal cortex resolves feature ambiguity in configural object recognition and perceptual oddity tasks. Learn Mem 14:821–832.

Bartra O, McGuire JT, Kable JW (2013) The valuation system: A coordinate-based meta-analysis of BOLD fMRI experiments examining neural correlates of subjective value. Neuroimage 76:412–427.

Basten U, Biele G, Heekeren HR, Fiebach CJ (2010) How the brain integrates costs and benefits during decision making. Proc Natl Acad Sci U S A 107:21767–21772.

Behzadi Y, Restom K, Liau J, Liu TT (2007) A Component Based Noise Correction Method (CompCor) for BOLD and Perfusion Based fMRI. Neuroimage 37:90–101.

Bettman JR, Luce MF, Payne JW (1998) Constructive Consumer Choice Processes. J Consum Res 25:187–217.

Botvinik-Nezer R, Salomon T, Schonberg T (2020) Enhanced Bottom-Up and Reduced Top-Down fMRI Activity Is Related to Long-Lasting Nonreinforced Behavioral Change. Cereb Cortex 30:858–874.

Brainard DH (1997) The Psychophysics Toolbox. Spat Vis 10:433–436.

Busemeyer JR, Gluth S, Rieskamp J, Turner BM (2019) Cognitive and Neural Bases of Multi-Attribute, Multi-Alternative, Value-based Decisions. Trends in Cognitive Sciences 23:251–263.

Bussey TJ, Saksida LM (2002) The organization of visual object representations: a connectionist model of effects of lesions in perirhinal cortex. Eur J Neurosci 15:355–364.

Bussey TJ, Saksida LM, Murray EA (2005) The perceptual-mnemonic/feature conjunction model of perirhinal cortex function. Q J Exp Psychol B 58:269–282.

Chib VS, Rangel A, Shimojo S, O’Doherty JP (2009) Evidence for a Common Representation of Decision Values for Dissimilar Goods in Human Ventromedial Prefrontal Cortex. J Neurosci 29:12315–12320.

Clark AM, Bouret S, Young AM, Murray EA, Richmond BJ (2013) Interaction Between Orbital Prefrontal and Rhinal Cortex Is Required for Normal Estimates of Expected Value. J Neurosci 33:1833–1845.

Cox RW, Hyde JS (1997) Software tools for analysis and visualization of fMRI data. NMR in Biomedicine 10:171–178.

Dale AM, Fischl B, Sereno MI (1999) Cortical surface-based analysis. I. Segmentation and surface reconstruction. Neuroimage 9:179–194.

Deichmann R, Gottfried JA, Hutton C, Turner R (2003) Optimized EPI for fMRI studies of the orbitofrontal cortex. Neuroimage 19:430–441.

Delgado MR, Beer JS, Fellows LK, Huettel SA, Platt ML, Quirk GJ, Schiller D (2016) Viewpoints: Dialogues on the functional role of the ventromedial prefrontal cortex. Nature Neuroscience 19:1545–1552.

Devlin JT, Price CJ (2007) Perirhinal contributions to human visual perception. Curr Biol 17:1484–1488.

Eldridge MAG, Lerchner W, Saunders RC, Kaneko H, Krausz KW, Gonzalez FJ, Ji B, Higuchi M, Minamimoto T, Richmond BJ (2016) Chemogenetic disconnection of monkey orbitofrontal and rhinal cortex reversibly disrupts reward value. Nat Neurosci 19:37–39.

Erez J, Cusack R, Kendall W, Barense MD (2016) Conjunctive Coding of Complex Object Features. Cereb Cortex 26:2271–2282.

Esteban O, Markiewicz CJ, Blair RW, Moodie CA, Isik AI, Erramuzpe A, Kent JD, Goncalves M, DuPre E, Snyder M, Oya H, Ghosh SS, Wright J, Durnez J, Poldrack RA, Gorgolewski KJ (2019) fMRIPrep: a robust preprocessing pipeline for functional MRI. Nature Methods 16:111.

Fonov V, Evans A, McKinstry R, Almli C, Collins D (2009) Unbiased nonlinear average age-appropriate brain templates from birth to adulthood. NeuroImage 47:S102.

Friston KJ, Williams S, Howard R, Frackowiak RS, Turner R (1996) Movement-related effects in fMRI time-series. Magn Reson Med 35:346–355.

Gitelman DR, Penny WD, Ashburner J, Friston KJ (2003) Modeling regional and psychophysiologic interactions in fMRI: the importance of hemodynamic deconvolution. Neuroimage 19:200–207.

Gorgolewski K, Burns CD, Madison C, Clark D, Halchenko YO, Waskom ML, Ghosh SS (2011) Nipype: a flexible, lightweight and extensible neuroimaging data processing framework in python. Front Neuroinform 5:13.

Gorgolewski KJ et al. (2016) The brain imaging data structure, a format for organizing and describing outputs of neuroimaging experiments. Sci Data 3:160044.

Greve DN, Fischl B (2009) Accurate and robust brain image alignment using boundary-based registration. Neuroimage 48:63–72.

Heide VD, JR, Skipper LM, Klobusicky E, Olson IR (2013) Dissecting the uncinate fasciculus: disorders, controversies and a hypothesis. Brain 136:1692–1707.

Hunt LT, Dolan RJ, Behrens TEJ (2014) Hierarchical competitions subserving multi-attribute choice. Nat Neurosci 17:1613–1622.

Jenkinson M, Bannister P, Brady M, Smith S (2002) Improved Optimization for the Robust and Accurate Linear Registration and Motion Correction of Brain Images. NeuroImage 17:825–841.

Kahnt T, Heinzle J, Park SQ, Haynes J-D (2011) Decoding different roles for vmPFC and dlPFC in multi-attribute decision making. Neuroimage 56:709–715.

Kaskan PM, Costa VD, Eaton HP, Zemskova JA, Mitz AR, Leopold DA, Ungerleider LG, Murray EA (2017) Learned Value Shapes Responses to Objects in Frontal and Ventral Stream Networks in Macaque Monkeys. Cereb Cortex 27:2739–2757.

Kennerley SW, Dahmubed AF, Lara AH, Wallis JD (2008) Neurons in the Frontal Lobe Encode the Value of Multiple Decision Variables. Journal of Cognitive Neuroscience 21:1162–1178.

Klein A, Ghosh SS, Bao FS, Giard J, Häme Y, Stavsky E, Lee N, Rossa B, Reuter M, Neto EC, Keshavan A (2017) Mindboggling morphometry of human brains. PLOS Computational Biology 13:e1005350.

Kondo H, Saleem KS, Price JL (2005) Differential connections of the perirhinal and parahippocampal cortex with the orbital and medial prefrontal networks in macaque monkeys. J Comp Neurol 493:479–509.

Krajbich I (2019) Accounting for attention in sequential sampling models of decision making. Current Opinion in Psychology 29:6–11.

Krajbich I, Armel C, Rangel A (2010) Visual fixations and the computation and comparison of value in simple choice. Nat Neurosci 13:1292–1298.

Kurtz-David V, Persitz D, Webb R, Levy DJ (2019) The neural computation of inconsistent choice behavior. Nature Communications 10:1583.

Lanczos C (1964) Evaluation of Noisy Data. Journal of the Society for Industrial and Applied Mathematics: Series B, Numerical Analysis 1:76–85.

Larsen T, O’Doherty JP (2014) Uncovering the spatio-temporal dynamics of value-based decision-making in the human brain: a combined fMRI–EEG study. Philos Trans R Soc Lond B Biol Sci 369 Available at: https://www.ncbi.nlm.nih.gov/pmc/articles/PMC4186227/ [Accessed August 7, 2020].

Levy DJ, Glimcher PW (2012) The root of all value: a neural common currency for choice. Curr Opin Neurobiol 22:1027–1038.

Lim S-L, O’Doherty JP, Rangel A (2013) Stimulus Value Signals in Ventromedial PFC Reflect the Integration of Attribute Value Signals Computed in Fusiform Gyrus and Posterior Superior Temporal Gyrus. J Neurosci 33:8729–8741.

McLaren DG, Ries ML, Xu G, Johnson SC (2012) A generalized form of context-dependent psychophysiological interactions (gPPI): a comparison to standard approaches. Neuroimage 61:1277–1286.

Mogami T, Tanaka K (2006) Reward Association Affects Neuronal Responses to Visual Stimuli in Macaque TE and Perirhinal Cortices. J Neurosci 26:6761–6770.

Mumford JA (2007) A Guide to Calculating Percent Change with Featquery. Available at: http://mumford.fmripower.org/perchange_guide.pdf.

Mumford JA, Nichols T (2008) Power Calculation for Group fMRI Studies Accounting for Arbitrary Design and Temporal Autocorrelation. Neuroimage 39:261–268.

Munafò MR, Smith GD (2018) Robust research needs many lines of evidence. Nature 553:399–401.

Murray EA, Bussey TJ, Saksida LM (2007) Visual Perception and Memory: A New View of Medial Temporal Lobe Function in Primates and Rodents. Annual Review of Neuroscience 30:99–122.

Nelissen K, Jarraya B, Arsenault JT, Rosen BR, Wald LL, Mandeville JB, Marota JJ, Vanduffel W (2012) Neural correlates of the formation and retention of cocaine-induced stimulus-reward associations. Biol Psychiatry 72:422–428.

Park SQ, Kahnt T, Rieskamp J, Heekeren HR (2011) Neurobiology of value integration: when value impacts valuation. J Neurosci 31:9307–9314.

Pelletier G, Fellows LK (2019) A Critical Role for Human Ventromedial Frontal Lobe in Value Comparison of Complex Objects Based on Attribute Configuration. J Neurosci 39:4124–4132.

Persichetti AS, Aguirre GK, Thompson-Schill SL (2015) Value is in the eye of the beholder: early visual cortex codes monetary value of objects during a diverted attention task. J Cogn Neurosci 27:893–901.

Philiastides MG, Biele G, Heekeren HR (2010) A mechanistic account of value computation in the human brain. Proc Natl Acad Sci U S A 107:9430–9435.

Power JD, Mitra A, Laumann TO, Snyder AZ, Schlaggar BL, Petersen SE (2014) Methods to detect, characterize, and remove motion artifact in resting state fMRI. Neuroimage 84:320–341.

Riesenhuber M, Poggio T (1999) Hierarchical models of object recognition in cortex. Nat Neurosci 2:1019–1025.

Rushworth MFS, Behrens TEJ (2008) Choice, uncertainty and value in prefrontal and cingulate cortex. Nat Neurosci 11:389–397.

Russo JE, Dosher BA (1983) Strategies for multiattribute binary choice. J Exp Psychol Learn Mem Cogn 9:676–696.

Salomon T, Botvinik-Nezer R, Oren S, Schonberg T (2020) Enhanced striatal and prefrontal activity is associated with individual differences in nonreinforced preference change for faces. Human Brain Mapping 41:1043–1060.

Schonberg T, Bakkour A, Hover AM, Mumford JA, Nagar L, Perez J, Poldrack RA (2014) Changing value through cued approach: an automatic mechanism of behavior change. Nature Neuroscience 17:625–630.

Schonberg T, Katz LN (2020) A Neural Pathway for Nonreinforced Preference Change. Trends in Cognitive Sciences 0 Available at: https://www.cell.com/trends/cognitive-sciences/abstract/S1364-6613(20)30104-2 [Accessed May 21, 2020].

Serences JT (2008) Value-Based Modulations in Human Visual Cortex. Neuron 60:1169–1181.

Shimojo S, Simion C, Shimojo E, Scheier C (2003) Gaze bias both reflects and influences preference. Nature Neuroscience 6:1317–1322.

Smith SM, Jenkinson M, Woolrich MW, Beckmann CF, Behrens TEJ, Johansen-Berg H, Bannister PR, De Luca M, Drobnjak I, Flitney DE, Niazy RK, Saunders J, Vickers J, Zhang Y, De Stefano N, Brady JM, Matthews PM (2004) Advances in functional and structural MR image analysis and implementation as FSL. Neuroimage 23 Suppl 1:S208–219.

Suzuki S, Cross L, O’Doherty JP (2017) Elucidating the underlying components of food valuation in the human orbitofrontal cortex. Nature Neuroscience 20:1780–1786.

Tustison NJ, Avants BB, Cook PA, Zheng Y, Egan A, Yushkevich PA, Gee JC (2010) N4ITK: Improved N3 Bias Correction. IEEE Transactions on Medical Imaging 29:1310–1320.

Vaidya AR, Fellows LK (2020) Under construction: ventral and lateral frontal lobe contributions to value-based decision-making and learning. F1000Res 9:158.

Vaidya AR, Sefranek M, Fellows LK (2018) Ventromedial Frontal Lobe Damage Alters how Specific Attributes are Weighed in Subjective Valuation. Cereb Cortex 28:3857–3867.

Wallis JD, Miller EK (2003) Neuronal activity in primate dorsolateral and orbital prefrontal cortex during performance of a reward preference task. Eur J Neurosci 18:2069–2081.

Watson HC, Wilding EL, Graham KS (2012) A Role for Perirhinal Cortex in Memory for Novel Object–Context Associations. J Neurosci 32:4473–4481.

Williams P (1998) Representational Organization of Multiple Exemplars of Object Categories. University of Massachusetts at Boston.

Xia C, Stolle D, Gidengil E, Fellows LK (2015) Lateral Orbitofrontal Cortex Links Social Impressions to Political Choices. J Neurosci 35:8507–8514.

Yoo SBM, Hayden BY (2018) Economic Choice as an Untangling of Options into Actions. Neuron 99:434–447.

Zhang Y, Brady M, Smith S (2001) Segmentation of brain MR images through a hidden Markov random field model and the expectation-maximization algorithm. IEEE Transactions on Medical Imaging 20:45–57.

